# Identification and implication of tissue-enriched ligands in epithelial-endothelial crosstalk during pancreas development

**DOI:** 10.1101/2022.04.19.488467

**Authors:** Manon Moulis, Steve Vincent Maurice Runser, Laura Glorieux, Nicolas Dauguet, Christophe Vanderaa, Laurent Gatto, Donatienne Tyteca, Patrick Henriet, Francesca M. Spagnoli, Dagmar Iber, Christophe E. Pierreux

## Abstract

Development of the pancreas is driven by an intrinsic program coordinated with signals from other cell types in the epithelial environment. These intercellular communications have been so far challenging to study because of the low concentration, localized production and diversity of the signals released. Here, we combined scRNAseq data with a computational interactomic approach to identify signals involved in the reciprocal interactions between the various cell types of the developing pancreas. This *in silico* approach yielded 40,607 potential ligand-target interactions between the different main pancreatic cell types. Among this vast network of interactions, we focused on three ligands potentially involved in communications between epithelial and endothelial cells. Bmp7 and Wnt7b, expressed by pancreatic epithelial cells and predicted to target endothelial cells, and Sema6d, involved in the reverse interaction. *In situ* hybridization confirmed the localized expression of *Bmp7* in the pancreatic epithelial tip cells and of *Wnt7b* in the trunk cells. On the contrary, *Sema6d* was enriched in endothelial cells. Functional experiments on *ex vivo* cultured pancreatic explants indicated that tip cell-produced Bmp7 restrained development of endothelial cells. This work identified ligands with a restricted tissular and cellular distribution and highlighted the role of Bmp7 in the intercellular communications shaping vessel development during pancreas organogenesis.

## INTRODUCTION

Organogenesis is a finely tuned process governed by the spatial, temporal and sequential expression of specific genes[1]. In every cell, control of gene expression is achieved by intrinsic and extrinsic, i.e. from the microenvironment, factors. Deciphering the catalogue of intrinsic and extrinsic factors, and understanding their connections, is a challenging but important task not only for fundamental knowledge but also to optimally orient differentiation of stem cells for tissue regeneration or engineering[2,3].

This is particularly relevant for the pancreas, an amphicrine gland that secretes digestive enzymes (exocrine function) and hormones regulating blood glucose homeostasis (endocrine function). Dysfunctional pancreas indeed causes major disorders such as diabetes or cancer, which remains important public health issues. Therefore, a better understanding of pancreas development and intercellular communications could improve differentiation protocols of hormone-producing cells or advance tissue engineering solution[4].

In mice, pancreas organogenesis starts around embryonic day (E) 8.5 when pancreatic progenitor cells expressing Pdx1 emerge from the foregut endoderm. These multipotent pancreatic progenitors proliferate and form the ventral and dorsal pancreatic buds. From E11.5, this 3D mass of non-polarized epithelial cells expands in the surrounding mesoderm-derived connective tissue and forms branches[5,6]. At the extremities or tip of these branches, epithelial cells express Ptf1a, Myc and Amylase; these tip cells will later enter the acinar differentiation program to give rise to the enzyme-producing acini. The more proximal or central part of the branches form a tubular plexus composed of trunk cells expressing Sox9 and Krt19. These trunk progenitors are bipotent and will later form the ducts transporting exocrine enzymes, as well as the endocrine islets of Langerhans. Along this differentiation program, epithelial cells are in close contact with mesenchymal cells and endothelial cells[7]. The mesenchyme is critical for pancreas development since its depletion alters epithelial morphogenesis and differentiation[8–11]. The endothelium also plays important roles in pancreas development. Signals from the endothelium are initially required for pancreatic budding[12] and later on for epithelial growth, endocrine and acinar differentiation[13–15]. Interestingly, at E11.5, endothelial cells are located all around the pancreatic bud, but from E13.5 they progressively and predominantly localize near the trunk cells at a distance from tip cells[14]. This blood vessel regionalization has been attributed to the preferential expression of Vegfa by the trunk cells[14], but other signals, e.g. preventing blood vessels localization around tip cells, must exist.

Recent studies have used transcriptomics to highlight the cellular heterogeneity[16,17], profile lineage dynamics[18] and decipher cell communication[19]. Here, we applied a computational interactomic analysis to create a repertoire/catalogue of potential intercellular communications in the developing pancreas, and to identify potential ligands involved in the reciprocal epithelial-endothelial crosstalk. To this aim, we combined single-cell RNAseq data[18] with the NicheNet framework[20]. We selected the endothelial ligand semaphorin 6d (Sema6d), and the epithelial ligands Wnt family member 7b (Wnt7b) and bone morphogenetic protein 7 (Bmp7) and validated their tissue localization by *in situ* hybridization. We then assessed the biological effects of Bmp7 on E12.5 pancreatic explants and demonstrated that localized Bmp7 production by the epithelial tip cells impairs development of blood vessels.

## RESULTS

### Cellular heterogeneity in the developing mouse pancreas

The gene expression profiles of E12.5 mouse pancreatic cells were obtained from a previously published single-cell RNAseq dataset[18]. Transcriptomic data analysis resulted in the separation of the cells in twelve clusters (Fig. 1A), which were then identified with known marker genes of specific cell types (Fig. 1B and Table S1). The epithelial cells (clusters 0-3) were easily picked out based on their expression of *Cdh1* and *Cldn6*[21,22]. Their abundance allowed us to distinctly identify four different epithelial subpopulations. The trunk cells (cluster 0) exhibited high expression levels of *Spp1* and *Sox9*[23,24]. The tip cells (cluster 1) evidently expressed tip marker genes such as *Ptf1a* and *Amy2b*[25,26]. Finally, two populations of endocrine cells (clusters 2 and 3) contained high levels of *Pax4*, insulin and glucagon transcripts, respectively[26,27]. Mesenchymal cells (clusters 4-6) expressed the marker *Col1a1*[28], and two subpopulations out of the three identified expressed the mesothelial markers *Wt1* and *Upk3b*[29,30]. We also found two immune subpopulations (clusters 8 and 9), and neuronal (cluster 10) and erythrocyte (cluster 11) progenitors, as already observed in mouse at later stages and in human[18,31,32]. A small subset of 226 cells (cluster 7) was identified as being endothelial based on their high expression levels of *Cdh5* and *Kdr*[33,34]. The low number of endothelial cells did not allow to readily identify subpopulations. However, reclustering of the isolated cells constituting cluster 7 allowed the identification of three endothelial subpopulations (Fig. 1C and 1D). The most abundant one (cluster A) shared markers of arterial and tip endothelial cells. The second one (cluster B), co-expressed venous and stalk cell markers, while the third one (cluster C) expressed lymphatic markers. Tip and trunk cells play fundamental role during angiogenesis. Endothelial tip cells migrate in response to pro-angiogenic factors, whereas stalk cells trail behind tip cells, proliferate and give rise to the future quiescent cells of mature vessels[35].

**Figure 1:**
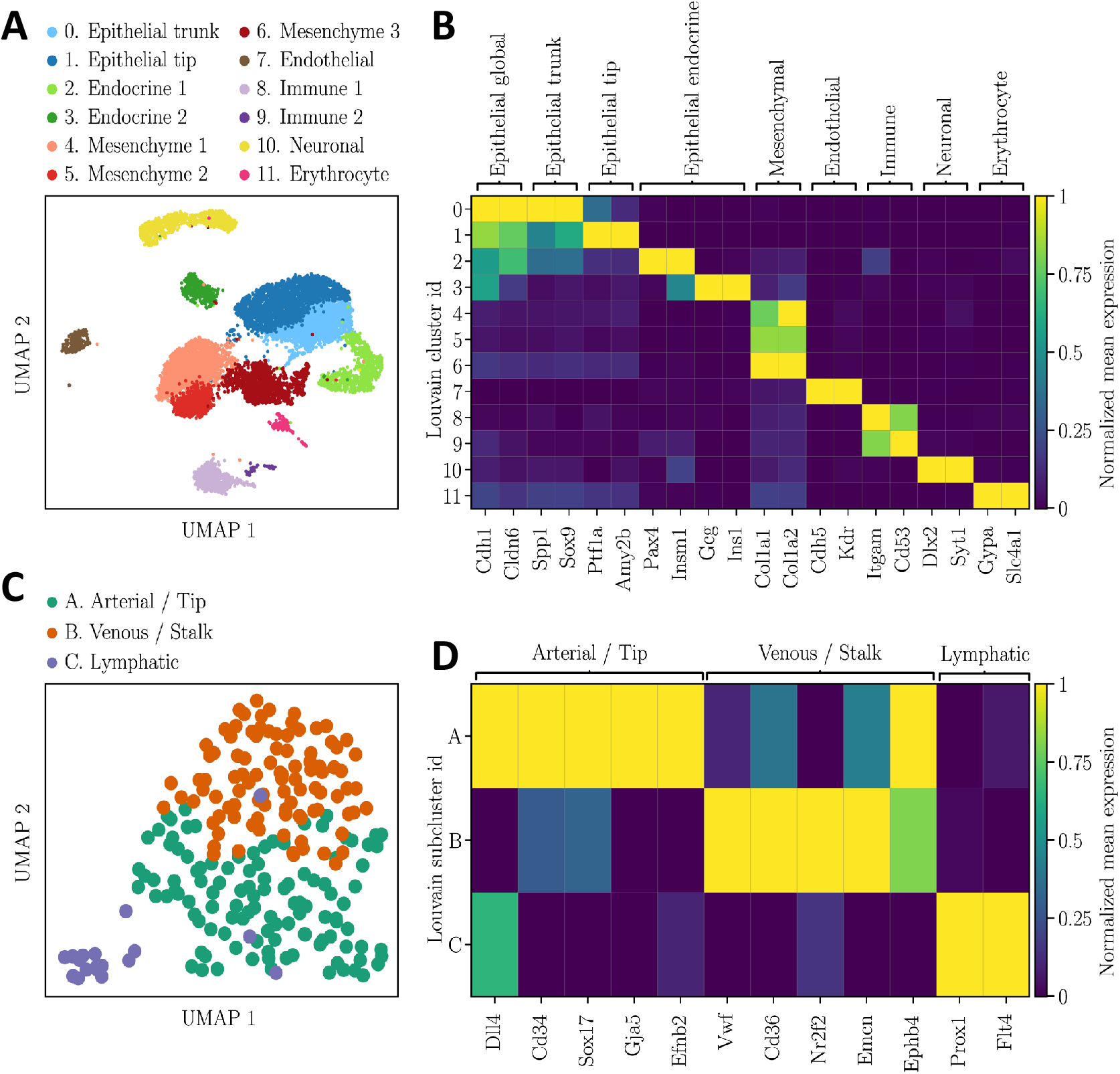
Identification of cellular subpopulations in murine E12.5 pancreas. (**A**) UMAP plot of the dataset E12.5 pancreatic cells colored according to their Louvain clusters. The result of the clusters manual annotation is given in legend. The cluster number 7 (endothelial cells) was reclustered with the Louvain algorithm with the aim to detect endothelial subpopulations. (**B**) Matrix plot of the Louvain clusters normalized mean expression levels of known marker genes. For each marker gene, the mean expression levels were normalized by first subtracting the minimum value and then dividing by the maximum value. (**C**) UMAP plot of the endothelial cells colored according to their Louvain subclusters. (**D**) Matrix plot of the endothelial subclusters normalized mean expression levels.

### Communications between the pancreatic cell populations and ligands of the epithelial-endothelial crosstalk

To facilitate our analysis of intercellular communications within the developing pancreas, we classified pancreatic cells in six major populations (epithelial, mesenchyme, endothelial, immune, neuronal and erythroblastic), and studied their reciprocal interactions. To do so, we used the NicheNet pipeline[20] to predict if ligands in a given cell population regulate the gene expression profiles in the five other cell populations. After ligand and target genes selection, NicheNet uses a built-in database of prior knowledge to infer how a set of ligands emitted by a sender population might regulate a set of target genes in a receiver population (Fig. 2A).

**Figure 2:**
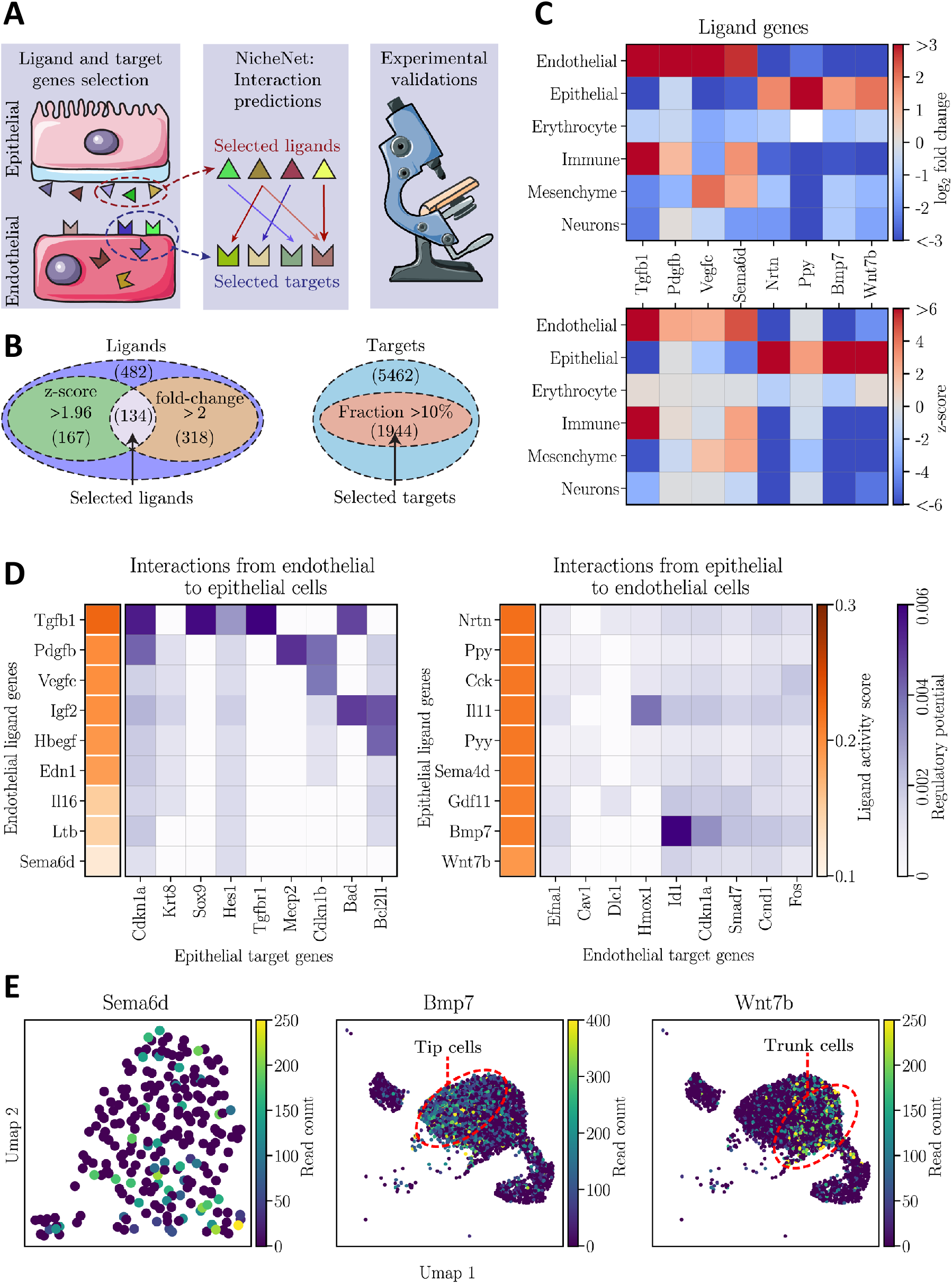
Interactome analysis using NicheNet database predicted active ligands potentially involved in pancreas vascular development and epithelial morphogenesis. (**A**) Schematic representation of the interactomic and experimental steps. The first step consisted in ligand and target genes selection in sender (*e*.*g*. epithelial) and receiver (*e*.*g*. endothelial) populations. Then, interactions between the selected ligands and targets were predicted with the NicheNet framework, allowing the prioritization of potential active ligands. Finally, some ligands were validated experimentally. (**B**) For a given population, ligand genes were considered expressed when they exhibited a z-score above 1.96 for the Wilcoxon signed-rank test and a fold change above 2. On the other hand, the target signaling genes were considered expressed in a population when at least a fraction of 10% of the cells constituting the population had 1 UMI (Unique Molecular Identifier) of the gene. The indicative numbers are detailed in the Material and Methods, section Interactomic analysis. (**C**) Matrix plots of the log_2_ fold change and z-score for the Wilcoxon signed-rank test of some endothelial and epithelial ligand genes which were selected based on the aforementioned thresholds. (**D**) NicheNet’s interactomic predictions from endothelial to epithelial (left panel) and from epithelial to endothelial (right panel). The ligands are ranked based on their activity scores (orange color map) while their regulation potential on target genes are colored in violet. (**E**) UMAP plots of the expression level of Sema6d in the endothelial cells as well as Bmp7 and Wnt7b in the epithelial cells.

In each sender population, we selected the ligand genes having a z-score above 1.96 for the Wilcoxon signed-rank test and a fold change above 2 to ascertain that the selected ligands would be later detectable on the stained tissue sections (Fig. 2B). In total, 134 ligands, out of the 482 ligands selected, met these two conditions in at least one population. In each receiver population, we selected as targets all the genes involved in a signal transduction pathway that were expressed by at least 10% of the population. We found 1944 signal transduction target genes that respected this 10% constrain in at least one of the major populations (Fig. 2B). An illustration of the fold change and z-score for some ligands is shown in Fig. 2C.

By combining the transcriptomic dataset with the NicheNet framework we identified 40,607 potential ligand-target interactions between the six major pancreatic cell populations (Table S2). This network of interactions being too large to be fully covered in this study, we focused on the interactions between the endothelial and epithelial cells. Among the predicted most active endothelial and epithelial ligands (Fig. 2D) we further investigated, through experimental validation, the effects of the endothelial ligand *Sema6d* as well as the epithelial ligands *Bmp7* and *Wnt7b*. This selection was based on the expression profiles of the ligand and the presence of the receptor in the receiver populations. For example, despite its high activity score, we did not select *Tgfb1* because immune cells also highly expressed this ligand (Fig. 2B). *Pdgfb, Vegfc* and *Nrtn* were excluded because their receptors were respectively absent from the pancreatic epithelium and endothelium. *Ppy* was highly expressed by epithelial cells, but not homogeneously distributed within epithelium as indicated by its lower z-score (Fig. 2B). In contrast, the ligands selected for experimental validation presented interesting expression profiles based on scRNAseq (Fig. 2E). *Sema6d* was found homogeneously expressed in the endothelial subpopulations, indicating a potential effect on the epithelium from any vessel subtype. On the contrary, *Bmp7* and *Wnt7b* exhibited localized expressions respectively in the tip and trunk epithelial cells, suggesting different potential effects on the vasculature[14].

### Spatio-temporal expression of the selected endothelial ligand, Sema6d

In order to corroborate the predictions of our interactomic analysis, we first verified the expression profiles of the three selected ligands (Wnt7b, Bmp7 and Sema6d). This was done by localizing the ligand-expressing cells on E10.5 to E14.5 pancreatic tissue sections through fluorescent *in situ* hybridization (RNAScope) coupled with immunolabeling of the epithelium (E-Cadherin) and endothelium (VE-Cadherin) (Fig. 3). Epithelial growth and branching morphogenesis were clearly illustrated on the low magnification images with the E-cadherin labelling (white in Fig. 3). Blood vessels, mostly peripheral at E10.5, progressively penetrated in between growing epithelial branches and contained some autofluorescent erythrocytes, indicating perfusion at E14.5 (green in Fig. 3).

**Figure 3:**
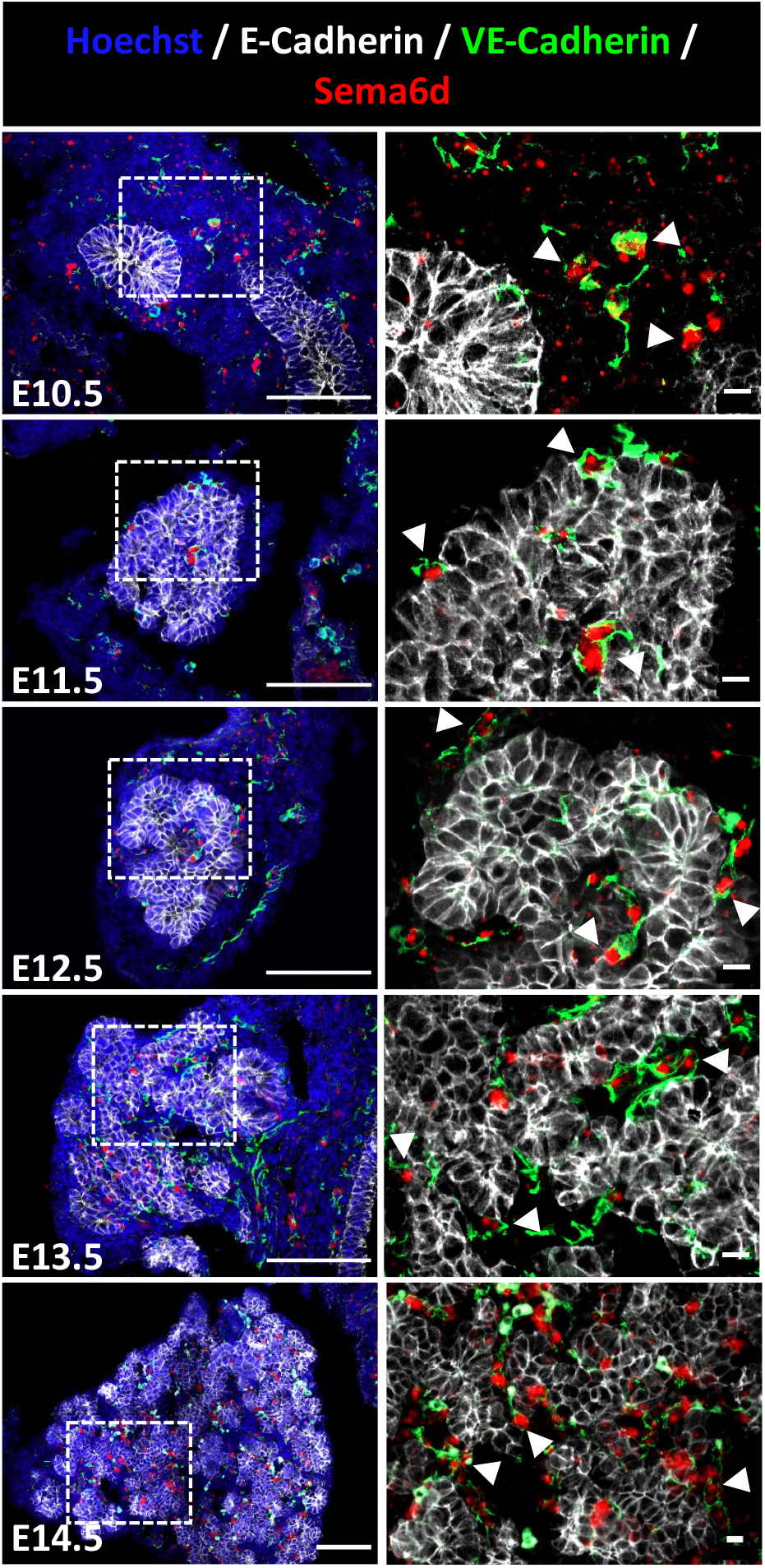
Localization of *Sema6d* expressing cells in developing pancreas. Detection of *Sema6d* transcripts by *in situ* hybridization (in red) on pancreatic tissue sections from E10.5 to E14.5, combined with an immunolabeling of E-Cadherin (white) and VE-Cadherin (green), to respectively detect the epithelium and vessels. Nuclei were counterstained with Hoechst (blue). Regions delineated by dashed lines on the left images are magnified on the right. Arrowheads indicate VE-Cadherin positive endothelial cells expressing *Sema6d* mRNA. Scale bars: 100 µm (at left) and 10 µm (at right).

In accordance with *in silico* analysis, *Sema6d* was predominantly found in the vascular compartment, showing a co-localization with VE-Cadherin positive endothelial cells (arrows in Fig. 3), but also in some circulating immune progenitors and mesenchymal cells. By RT-qPCR, abundance of *Sema6d* mRNA expression level remained stable between E11.5 and E15.5 (Fig. S1 A), and similar to that found in other endodermal (stomach, intestine, liver and lung) and mesodermal (heart and spleen) organs at E15.5 (Fig. S1 B). Even when significant differences in the expression levels of *Sema6d* were detected between the organs, the fold change was smaller than a factor of 4. We therefore confirmed the expression of Sema6d in endothelial cells during pancreas development and morphogenesis. Investigation of the potential regulatory role of Sema6d on pancreas development and morphogenesis should take into account that this ligand is non-secreted and known to act in a juxtacrine manner[36].

### Validation of the expression pattern of the epithelial ligands, Wnt7b and Bmp7

NicheNet analysis highlighted two epithelial ligands that could signal towards the vascular compartment, Wnt7b and Bmp7, as already reported[37,38]. In addition, *in silico* analysis revealed that these ligands were produced by two different epithelial cell populations of the developing pancreas, namely the trunk cells for Wnt7b and tip cells for Bmp7 (Fig. 2E). *In situ* hybridization experiments confirmed this prediction with a distinct trunk sublocalization for *Wnt7b*, and an increased abundance of *Bmp7* in the tip cells at the periphery (Fig. 4A and 4B). We observed that *Wnt7b* expression was homogeneously distributed in the pancreatic epithelium, and excluded from duodenum at E10.5, whereas *Bmp7* expression was already enriched at the periphery at this stage. The trunk- and tip-enriched expression patterns became more obvious with pancreas development, i.e. at E13.5 and E14.5 with ductal epithelium expressing *Wnt7b* and acinar buds expressing *Bmp7*. Based on the *in situ* hybridization experiments, we concluded that the expression of these two ligands is spatially restricted but maintained during the developmental stages analyzed, thereby suggesting a prolonged biological effect during pancreas development.

**Figure 4:**
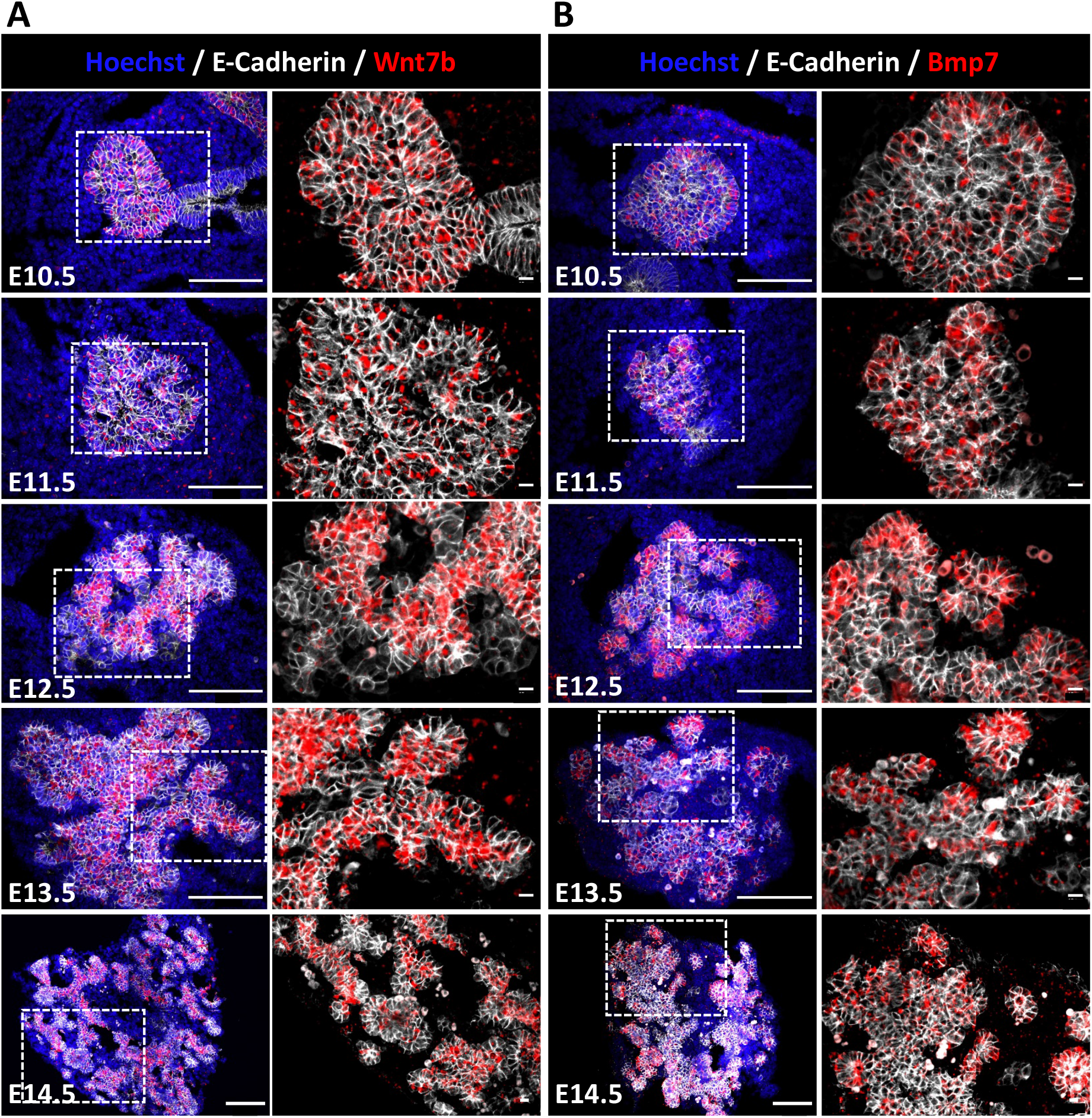
Localization of *Wnt7b* and *Bmp7* expressing cells in developing pancreas. Detection of *Wnt7b* (**A**) and *Bmp7* (**B**) transcripts by *in situ* hybridization (in red) on pancreatic tissue sections from E10.5 to E14.5, combined with an immunolabeling of E-Cadherin (white) to detect the epithelium. Nuclei were counterstained with Hoechst (blue). Regions delineated by dashed lines on the left images are magnified on the right, highlighting *Wnt7b* and *Bmp7* mRNA expression in E-Cadherin positive trunk and tip cells, respectively. Scale bars: 100 µm (at left) and 10 µm (at right).

After validation of *Wnt7b* and *Bmp7* localization, we quantified their expression level by RT-qPCR (Fig. S2 A). Since these ligands are expressed by epithelial cells, we normalized their expression levels to that of *cadherin-1* (*Cdh1*)/E-cadherin, to account for epithelial proliferation and expansion (Fig. 4A and 4B, left panels). The relative expression level of *Wnt7b* and *Bmp7* was stable from E11.5 to E13.5 but showed a significant decrease at E15.5, supporting the progressive regionalization of these two ligands within the pancreatic epithelium. Expression of *Wnt7b* and *Bmp7* was not only interesting for the regionalized and persistent (E10.5 to E15.5) characteristics but also for the higher level in the pancreas as compared to other organs (Fig. S2 B and C). Indeed, we found that pancreatic expression of *Wnt7b* and *Bmp7* was 4 to 50 times more important than in other organs, suggesting a particular role for these ligands in the pancreas.

### Biological effect of the epithelial tip-enriched ligand Bmp7 on pancreas development

Finally, we decided to test whether the Bmp7 epithelial ligand could impact on pancreas development and more specifically on the vascular compartment. We used microdissected E12.5 pancreatic explants cultured on a filter for up to 72 hours. This *ex vivo* culture system has been widely used since it reproduces pancreatic differentiation and morphogenesis[14,39–41], and is suitable to test the effect of soluble proteins such as Bmp[42].

We first analysed the Bmp responsiveness of E12.5 pancreas, by incubating microdissected pancreas with a Bmp7 recombinant protein (BMP7: 400 ng/ml) for 90 minutes (Fig. S3 A). Pancreata were fixed and sections labelled with an antibody against the phosphorylated form of Smad 1/5, an indicator of effective Bmp signal transduction from the membrane to the nucleus[42]. We found nuclei positive for phospho Smad1/5 in untreated explants, probably due to the presence of endogenous Bmps, but more nuclei were labelled in BMP7-treated explants, indicating activation of the pathway (Fig. S3 A). Interestingly, we found that the phosphorylated form of Smad1/5 was visible in different cell types, including VE-Cadherin positive endothelial cells (Fig. S3 A). Pancreatic explants were then cultured with BMP7 or DMH-1 (3 µM), a selective inhibitor of the Bmp type-I receptor subtype Alk2 on which the Bmp7 ligand binds[43], and we measured the expression of Bmp target genes, *Id1, Id2* and *Id3*, by RT-qPCR after 48h of culture (Fig. 5A). Expression of these three genes was upregulated upon BMP7 treatment, and downregulated in DMH-1-treated explants, as compared to control explants. Furthermore, similar transcriptional effects were observed with primary culture of endothelial cells, confirming that endothelial cells can respond to BMP7 (Fig. S3 B). Macroscopically, expansion and morphogenesis of the pancreatic translucid epithelium were not affected by the treatments, even if BMP7-treated explants appeared smaller on the filter (Fig. S3 C). Histological analysis confirmed normal morphogenesis and development of the pancreatic epithelium (E-Cadherin in Fig. 5B and 5E), as well as the normal thickness (Hoechst in Fig. 5B, 5D, 5E and S3 E). In line with these observations, expression of *Cdh1* was not affected by BMP7 or DMH-1 treatments (Fig. S3 D). Interestingly, we found that BMP7-induced signalling affected the endothelial compartment, as reflected by the reduced VE-Cadherin labelling and quantification (Fig. 5B). To confirm that the decrease of VE-Cadherin surface is due to a loss of the vascular compartment and not to an effect on the sole expression of VE-Cadherin, we analysed other endothelial markers by RT-qPCR and immunolabeling. We first measured the expression level of *Pecam1* and *Cdh5* (VE-Cadherin) using the same extracts as for the *Id* genes. We found a decreased expression for these two endothelial markers in the presence of BMP7, but no effect of DMH-1 (Fig. 5C). Furthermore, the negative effect of BMP7 on pancreatic vasculature was also evidenced using a third vascular marker, endomucin (Fig. 5D). On the contrary, DMH-1 treatment did not increase vessels abundance (Fig. 5B-D). Lastly, co-labeling of endomucin with phospho Smad 1/5 indicated that the response to BMP7 was sustained since we detected phospho Smad 1/5 signals in explants treated for 72h in culture (Fig. 5D). Phospho Smad 1/5 was found in peripheral mesenchymal cells but also in endothelial cells (insets in Fig. 5D). Altogether, these results suggest that BMP7 is able to activate Smad1/5 signalling in endothelial cells and that this activation negatively impact blood vessels abundance.

**Figure 5:**
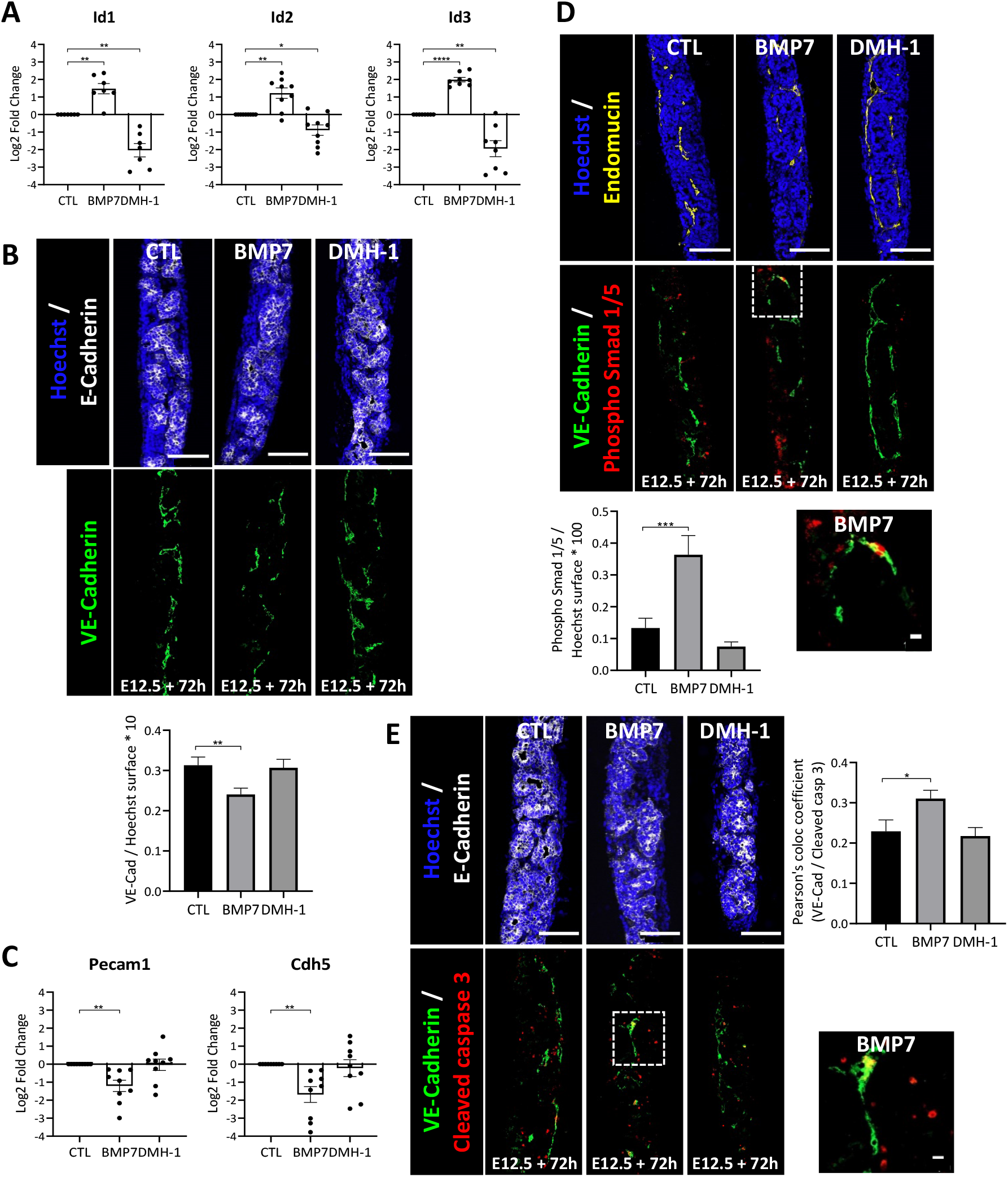
Decreased vascular density, via increased endothelial cell apoptosis, upon Bmp7 treatment of pancreatic explants. Pancreatic explants were treated with Bmp7 recombinant protein (BMP7, 400 ng/mL), Bmp signalling inhibitor DMH-1 (3 µM), or left untreated (CTL), for 48h (**A, C**) or 72h (**B, D, E**). (**A**) RT-qPCR analysis of Bmp target genes *Id1, Id2* and *Id3* normalized to *Actb* and *Rpl27*, and presented in log2 fold change (n=7-9). Expression of *Id* genes was increased by BMP7 and decreased by DMH-1. (**B**) Immunolabeling of epithelial E-Cadherin (white) and endothelial VE-Cadherin (green) cells, with Hoechst nuclei counterstaining (blue). Below, quantification of VE-Cadherin-labelled surface reported to total Hoechst^+^ surface (n=7) showed decreased vascular density with BMP7. Scale bars: 100 µm. (**C**) RT-qPCR analysis of endothelial (*Pecam1* and *Cdh5*) markers normalized to *Actb* and *Rpl27*, and presented in log2 fold change (n=9). Expression of both endothelial markers was decreased by BMP7. (**D**) Immunolabeling of phospho Smad 1/5 (red) and endothelial endomucin (yellow) or VE-Cadherin (green), with Hoechst nuclei counterstaining (blue). At right, quantification of phospho Smad 1/5^+^ surface reported to total Hoechst+ surface (n=4) showed increased phospho Smad 1/5^+^ surface in BMP7-treated explants. Region delineated by dashed lines on the BMP7 explant is magnified below, and highlights an endothelial cell with a phospho Smad 1/5^+^ nucleus. Scale bars: 100 µm and 10 µm (inset). (**E**) Immunolabeling of cleaved caspase 3 (red), epithelial E-Cadherin (white) and endothelial VE-Cadherin (green) cells, with Hoechst nuclei counterstaining (blue). Measure of the Pearson’s correlation coefficient for colocalization between cleaved caspase 3- and VE-Cadherin-pixels (n=4) showed increased VE-Cadherin/cleaved caspase 3 colocalization in BMP7-treated explants. Region delineated by dashed lines in the BMP7 condition is magnified below, and illustrates endothelial cell apoptosis. Scale bars: 100 µm and 10 µm (inset). One-way ANOVA (comparison to CTL): * p<0.05, ** p<0.005, *** p<0.0005, **** p<0.0001.

We then tested the hypothesis that the inhibitory effect of Bmp7 ligand on vessels could be due to increased apoptosis. Explants were treated with BMP7 or DMH-1 and then labelled with an antibody against cleaved caspase 3 (Fig. 5E). Overall apoptosis did not vary significantly between the three conditions (data not shown). However, we found that apoptosis of endothelial cells, visualized by colocalization of VE-Cadherin with cleaved caspase 3, was significantly higher in BMP7-treated explants, as compared to control explants, thereby suggesting a BMP7-induced endothelial-specific apoptosis. Finally, since we previously demonstrated that blood vessels density and localization control acinar differentiation[14,41], we tested whether addition of exogenous Bmp7 ligand could impact on pancreatic acinar differentiation through the vasculature. Although not significant, BMP7 increased the expression of two acinar genes, namely *Amy2a* and *Ptf1a* (Fig. S3 D), while *Krt19* and *Sox9*, two ductal markers did not vary. Since phospho Smad1/5 was not observed in the pancreatic epithelium, we excluded an autocrine effect of BMP7, and propose a model in which Bmp7 ligand secretion by pancreatic tip cells would prevent endothelial cell expansion in the tip niche, thereby promoting acinar differentiation.

## DISCUSSION

In this study, we obtained the gene expression profiles of the six main pancreatic populations by analyzing a previously published E12.5 mouse single-cell RNAseq dataset[18]. We determined the ligands and signal transduction target genes expressed in each population, and predicted potential interactions between these ligands and target genes with the NicheNet framework[20]. This interactomic analysis yielded 40,607 potential ligand-target interactions between the main pancreatic populations. For practical reasons we limited our analysis and validation to the communications between epithelial and endothelial cells. Among the predicted ligands we investigated the endothelial ligand Semad6d and the epithelial ligands Bmp7 and Wnt7b. Through immunolocalization on pancreatic tissue sections, we confirmed the predicted localization of Sema6d in endothelial cells, and of Bmp7 and Wnt7b in the epithelial tip and trunk cells, respectively. Finally, using a 3D *ex vivo* culture system of the pancreas, we demonstrated an inhibitory effect of Bmp7 on the blood vessels development.

Our interactomic study was carried out between the global pancreatic populations, rather than its subpopulations, to identify important signals conserved at the population level. However, this approach can be refined for more specific biological questions. Firstly, by performing the interactomic analysis between the subpopulations; e.g. endocrine epithelium towards endothelium without the lymphatic subpopulation for precise research of endocrine ligands influencing blood vessels angiogenesis in developing islets. Secondly, by using other selection criteria for the ligand and target genes. For instance, we could have selected as targets only the genes modulated during angiogenesis for an oriented research on this topic. Regarding the interactomic analysis, it is also important to consider the biases introduced by the NicheNet database. Indeed, this framework integrates prior knowledge and, obviously, results depend on reported network information rather than on the cellular gene expression profiles in the dataset[44]. Thus, direct or indirect links between populations of interest could be inferred without functional relevance, but because they were described in other contexts. To obtain a proof of concept that ligands unveiled in the interactomic analysis are biologically functional, we followed an oriented approach. We focused on epithelial-endothelial reciprocal communications and selected three ligands displaying a clear *in silico* enrichment in a cell population, and having known (Wnt7b) or unknown (Sema6d and Bmp7) effects on pancreas development.

Although Sema6d effects have recently been studied in different contexts[36,45–47], the cell types expressing Sema6d are rarely specified. Using *in situ* hybridization of *Sema6d*, we confirmed the *in silico* analysis and described for the first time Sema6d expression in the developing mouse embryonic pancreas. Sema6d is restricted to the endothelium and its expression was maintained in this compartment from E11.5 to E15.5. In addition, we found that Semad6d endothelial-specific expression level was similar in different developing organs, thereby suggesting a similar endothelial expression pattern in these organs.

Expression and role of Wnt7b in the pancreatic epithelium have been described[48,49], but we here refined its tissue distribution and showed that it progressively became enriched in epithelial trunk cells, thereby validating scRNAseq data. The role of Wnt7b has been studied *in vivo* and shown to be important for progenitor growth at the expense of differentiation[48]. These authors also observed that overexpression of Wnt7b induces a disproportionate increase in mesenchyme. Unfortunately, blood vessels were not studied in these loss- and gain-of Wnt7b function[48]. Based on the trunk-enrichment of Wnt7b described in this study, it would be interesting to study Wnt7b role on the bipotent pancreatic trunk progenitors and to study the effect on blood vessels *in vivo* or in co-culture experiments[49]. Indeed, given the interactomic data revealing potential interaction of Wnt7b with endothelial cells and given its enriched expression by the trunk epithelium, like the pro-angiogenic factor Vegfa[14], one could propose that Wnt7b favors vessel recruitment and maintenance around the trunk epithelium, as suggested in the choroid and in cancer[37,50].

Bmp signalling, including Bmp7, has been studied in developing pancreas and shown to affect primarily the mesenchyme and indirectly pancreas development[51]. Our interactomic data and *in situ* hybridization studies support the epithelial origin of Bmp7. Furthermore, scRNAseq and *in situ* hybridization revealed an enrichment of this ligand in the epithelial tip cell population of the pancreas, thereby suggesting a local, and not global, role on the surrounding microenvironment. Indeed, NicheNet suggested an interaction of Bmp7 with the endothelial compartment of the stroma. This hypothesis was tested in explants and revealed increased apoptosis in endothelial cells. In this *ex vivo* cultured system, the inhibitory effect of Bmp7 on blood vessels was global since the whole pancreatic explants were incubated with exogenous Bmp7. In addition, since we detected phosphorylated Smad1/5 in mesenchymal cells in response to exogenous Bmp7, we cannot exclude an indirect effect on endothelial cells via the mesenchyme. However, interactomic data, localized expression pattern of Bmp7, and direct effect of Bmp7 on cultured endothelial cells, suggest a local and direct inhibitory effect on blood vessels. This is compatible with the work of Tate *et al*. who demonstrated that Bmp7 treatment of cultured endothelial cells caused a decrease in the expression of Vegfr2 and Fgfr1 receptors, in endothelial cell migration and tube formation, but also a decrease in tumor vessels density *in vivo* after treatment with a recombinant protein for Bmp7, attesting of an anti-angiogenic effect[38]. The inhibitory role of Bmp7 on vascular development is also compatible with the predominant localization of blood vessels around the trunk epithelial cells and their scarcity around pancreatic tip cells[7,14]. Altogether, one can propose that the localized production of the blood vessel inhibitory ligand, Bmp7, around tip cells could work in concert, but in an opposite manner, with the localized production of angiogenic Vegfa by the trunk cells. This hypothesis could be tested *in vivo* with transgenic localized overexpression, as already performed for Vegfa[14], or in engineered tissue. Surprisingly, and although Bmp signaling inhibition with DMH-1 decreased the expression of the *Id* Bmp target genes, DMH-1 had no effect on the expression of vascular markers, as measured by RT-qPCR or immunolabeling and did not increased vessel density in the explants. This could be explained by local inhibitory signal in a niche domain. Around the trunk, there is no Bmp7 produced and vessels are abundant, survive and proliferate due to the local action of Vegfa. Adding DMH-1 in the absence of inhibitory Bmp7 will have no effect in this niche. Around the tip epithelial cells, blood vessels are scarce probably due to the local production of inhibitory Bmp7 combined with the absence of the pro-angiogenic Vegfa. In this niche, DMH-1 will block the inhibitory Bmp7 signaling but the scarcity of blood vessels and the absence of pro-angiogenic signal will prevent detection of any effect in this *ex vivo* culture system.

This work thus unravels potential cellular interactions during pancreas development via an interactomic approach, transposable to other contexts (species, organs, pathologies, etc.). In addition, it provides a proof-of-concept for the discovery of new ligands potentially regulating intercellular communications during pancreas development. Specifically, we identified Bmp7 as a tip-cell enriched ligand that can prevent blood vessels expansion around the pancreatic tip niche, thereby adding a functional link shaping development of the pancreatic epithelium and endothelium.

## MATERIAL AND METHODS

### Data origin

The gene expression profiles of E12.5 mouse pancreatic cells were obtained from the previously published dataset GEO GSM3140915[18]. This dataset was particularly adapted to the current study as it was depleted from mesenchymal cells and thereby enriched in epithelial and endothelial cells. The raw single-cell data were demultiplexed and converted to FASTQ files with the 10x Genomics Cell Ranger pipeline (v6.1.2). The reads were mapped onto the mouse reference genome GRCm38 with the genome aligner STAR (v2.7.2a)[52]. The options used to parameterize the aligner are given with the available code. The expression levels of 50,686 genes were thus measured for 16,286 pancreatic cells.

### Data processing

The generated feature-barcode matrix was analyzed with the python Scanpy pipeline (v1.8.2)[52]. As a first preprocessing step, the cells with less than 1,500 UMIs (Unique Molecular Identifiers) or less than 2,000 active genes were filtered out from the dataset. As a second step, the fraction of mitochondrial versus cytoplasmic RNA was measured in each cell. A fraction higher than 0.05 is indicative of broken cells[53]. Fortunately, no cell in the dataset exhibited such a high level of mitochondrial RNA. A total of 10,822 cells were thus retained for further analysis.

To compare the expression level of each gene across different cells, we normalized the counts with respect to the library sizes (counts per million) and logarithmized them. To identify the highly variable genes (HVG), we used the approach based on the counts coefficients of variations developed by Satija *et al*[54]. 4,696 genes were thus identified as having highly variable expression levels across the cells of the dataset. To prevent unwanted source of variation among the cells, we regressed out the biological effects caused by the cell cycle and the mitochondrial genes expression. In addition, we regressed out the technical effect created by the sequencing depth. The resulting expression levels were then standardized such that each highly variable gene had a null mean expression and a unit variance across the cells.

### Clustering

The high-dimensional expression profile of each cell’s HVG was mapped to a lower dimensional space by computing the first 40 principal components. A neighborhood graph of the cells was constructed in this low dimensional space and then divided in clusters with the Louvain algorithm[55]. The obtained results were visualized on a UMAP plot[56]. The clusters were manually annotated based on known cell population marker genes (Table 1).

**Table 1:**
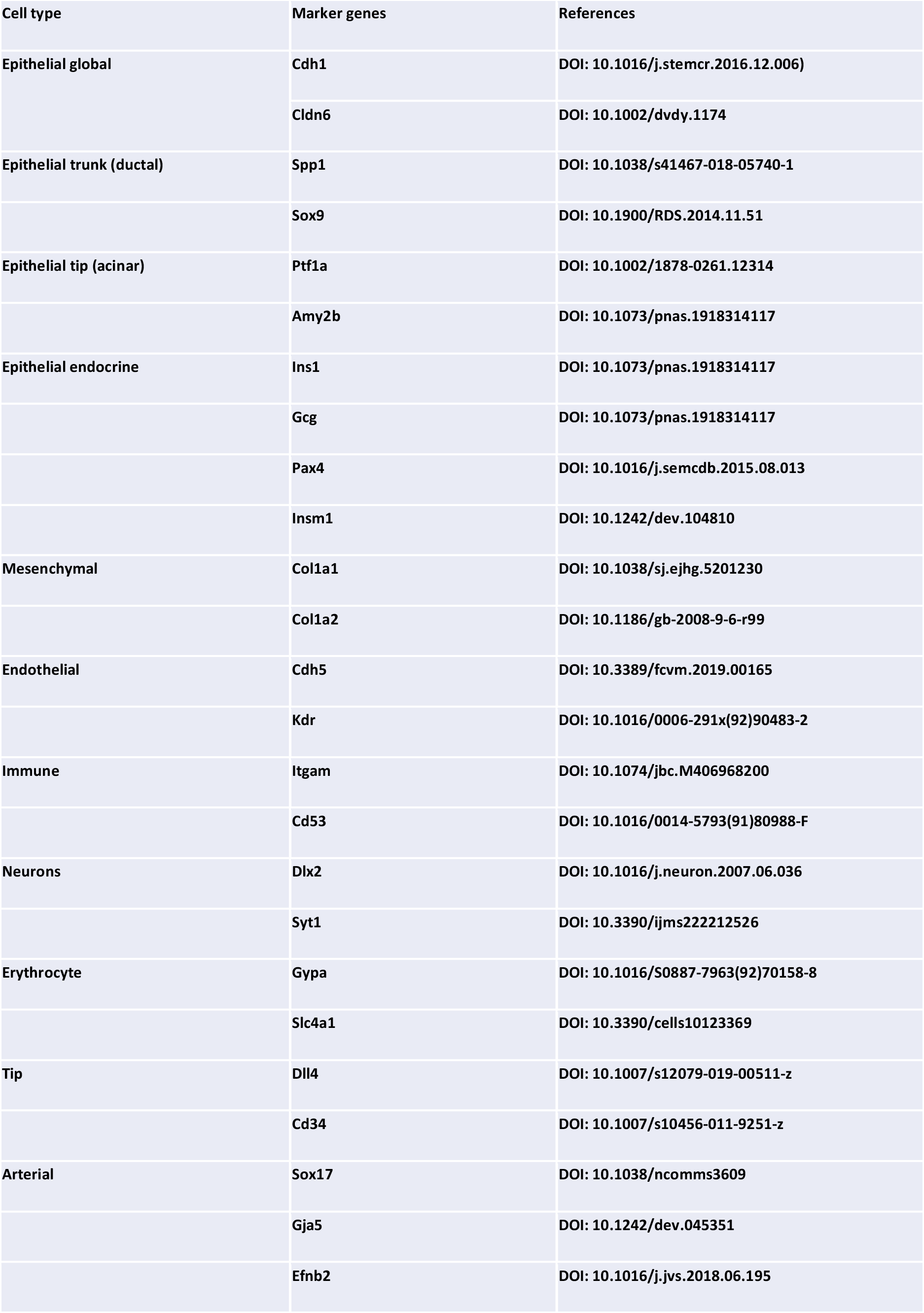
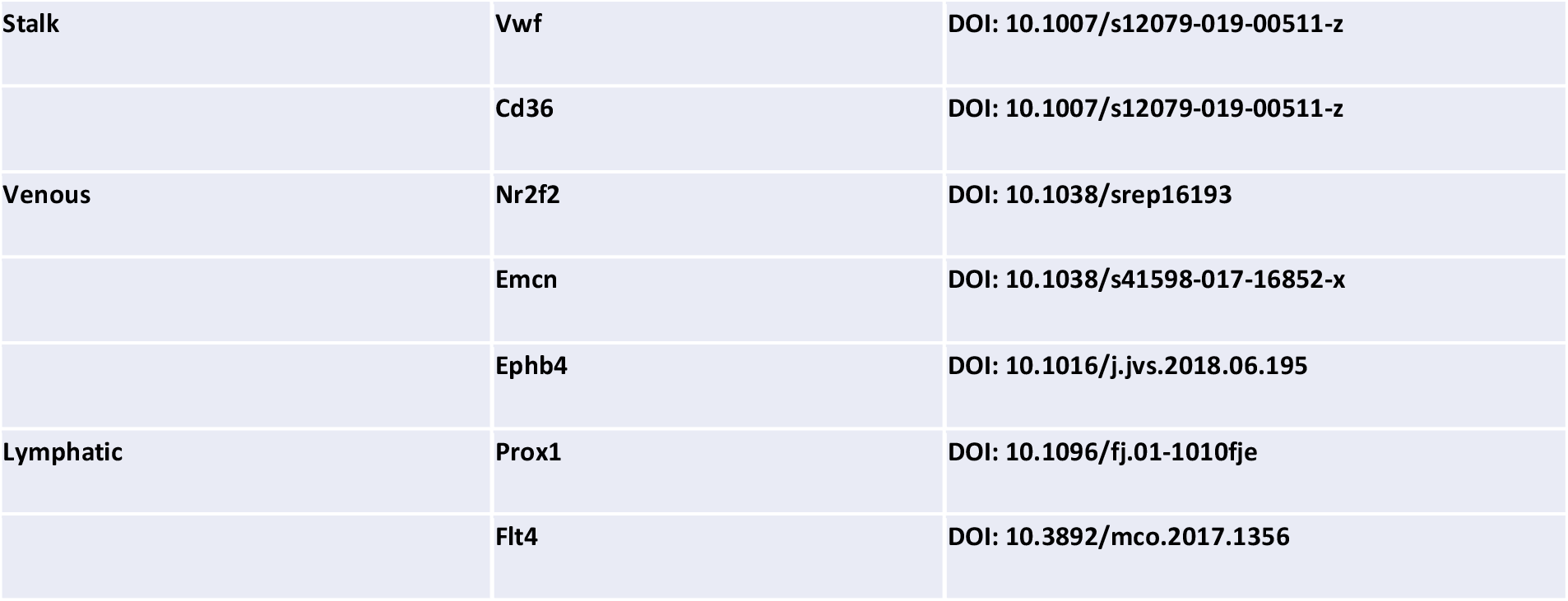
Marker gene references.

### Interactomic analysis

We employed the NicheNet framework[20] to unravel the network of cellular communications between the different general cell types detected. This framework predicts how the expression of certain target genes in a receiver population are affected by the expression of ligands in a sender population. We used the NicheNet pipeline on all the 36 possible combinations of sender/receiver populations. As a prerequisite, the model has to be made aware of the ligand and target genes expressed respectively in the sender and receiver populations.

The ligand genes were selected based on the annotations available on the Gene Ontology database (GO:00048018). We compared the expression distribution of each of these ligand genes in a given population against the other populations with the non-parametric Wilcoxon test. The ligands were considered significantly differentially expressed in a population when their z-scores exceeded 1.96. Out of 482 available ligand genes, 167 fulfilled this criterion in at least one of the populations. In addition, we computed the ratio between the average expression of a ligand gene in a given population and the average expression of the same ligand gene in all the other populations. This ratio that we name hereafter fold change was measured to ensure that the selected ligands would be clearly localizable in their respective populations during the *in situ* hybridization experiment. We considered a ligand to be overexpressed in a population when its fold change exceeded 2. Out of 482 available ligand genes, 318 were thus considered overexpressed. For the rest of the interactomic analysis, we only retained the 134 ligand genes that were significantly differentially expressed and overexpressed at the same time in at least one of the populations.

For the purpose of this analysis, all the genes having been identified as part of a signal transduction pathway (GO:0007165) were considered target genes. We deemed a target gene as being expressed in a population when at least 10% of the cells constituting the population had at least one UMI of the gene. 1944 genes fulfilled this criterion out of the 5462 available signaling genes.

### Data availability

The whole code used for the analysis as well as the figures 1 and 2 can be found at gitlab (URL: https://u.ethz.ch/mGSpS). The whole analysis pipeline combining the code with the feature-barcode matrix and the NicheNet interactomic predictions can be found at openbis (URL: https://u.ethz.ch/fYbuR).

### Animals and embryo dissection

Wild-type C57BL/6 mice (Jackson Laboratory) were raised and treated according to the NIH Guide for Care and Use of Laboratory Animals. Experiments were approved by the University Animal Ethical Committee, UCLouvain (2016/UCL/MD/005 and 2020/UCL/MD/011), and followed the recommendations of the ARRIVE guidelines. Males and females were mated, and the day of the vaginal plug was considered as embryonic day (E) 0.5. Pregnant females were sacrificed by cervical dislocation at the desired time point, and embryos were collected for further microdissection.

### RNAScope *in situ* hybridization assay coupled with immunofluorescence on paraffin sections

Tissue samples (E10.5: entire embryo, E11.5: abdomen, E12.5-E13.5-E14.5: stomach with pancreas) were fixed in 4% paraformaldehyde for 24h, embedded in paraffin using the Tissue-Tek VIP 6 (Sakura) tissue processor and of 6 µm sections obtained with a microtome (HM355S, Thermo Scientific). Z-shaped probes for Sema6d (565871), Wnt7b (401131), Bmp7 (407901), DapB (310043, negative control) and Ppib (313911, positive control) were hybridized on sections for 2h at 40°C in the HybEZ II oven as described[41]. Tissues were then blocked and immunolabeled as described in the section “Immunofluorescence on gelatin sections”. Slides were mounted and scanned with the Pannoramic P250 Digital Slide Scanner (3DHistech), and acquired with the Cell Observer Spinning Disk Confocal Microscope (Zeiss).

### Pancreatic explant culture and treatment

After microdissection and three washes in culture medium, E12.5 pancreatic dorsal buds were placed on microporous membranes (PICM01250, Millipore) at the air-medium interface. DMEM/F-12 medium (11039-021, Gibco) with 10% serum, 100 U/mL penicillin and 100 µg/mL streptomycin was further supplemented with 400 ng/mL of BMP7 recombinant protein (5666-BP, R&DSystems), 3 µM of DMH-1 (4126/10, R&DSystems), or vehicles. Explants were cultured for 48h or 72h, with medium renewal every day, and 10 µL of the culture medium was added on the explants on top of the filter 3 times per day.

### Immunofluorescence on gelatin sections and quantification

Pancreatic explants were fixed in 4% paraformaldehyde for 30 min, followed by equilibration in PBS-20% sucrose solution, embedding in PBS-15% sucrose-7.5% gelatin, and cryosectioning. Sections of 8 µm were obtained with a cryostat (CryoStar NX70, Thermo Scientific) and immersed for gelatin removal and antigen retrieval in citrate buffer (10 mM, pH 6) heated (microwave 750 W) 2 × 5 min. Sections were permeabilized 5 min with PBS-0.3% Triton X-100, blocked 45 min with PBS/0.3% Triton X-100/10% BSA/3% milk (blocking solution), and incubated on night at 4°C with primary antibodies (Table 2) diluted in blocking solution. After three washes with PBS/0.1% Triton X-100, sections were incubated with secondary fluorescent antibodies (AlexaFluor, Invitrogen) and Hoechst 33258 fluorescent nuclear dye (Sigma) in PBS/0.3% Triton X-100/10% BSA for 1h at room temperature, before 3 washes.

**Table 2:**
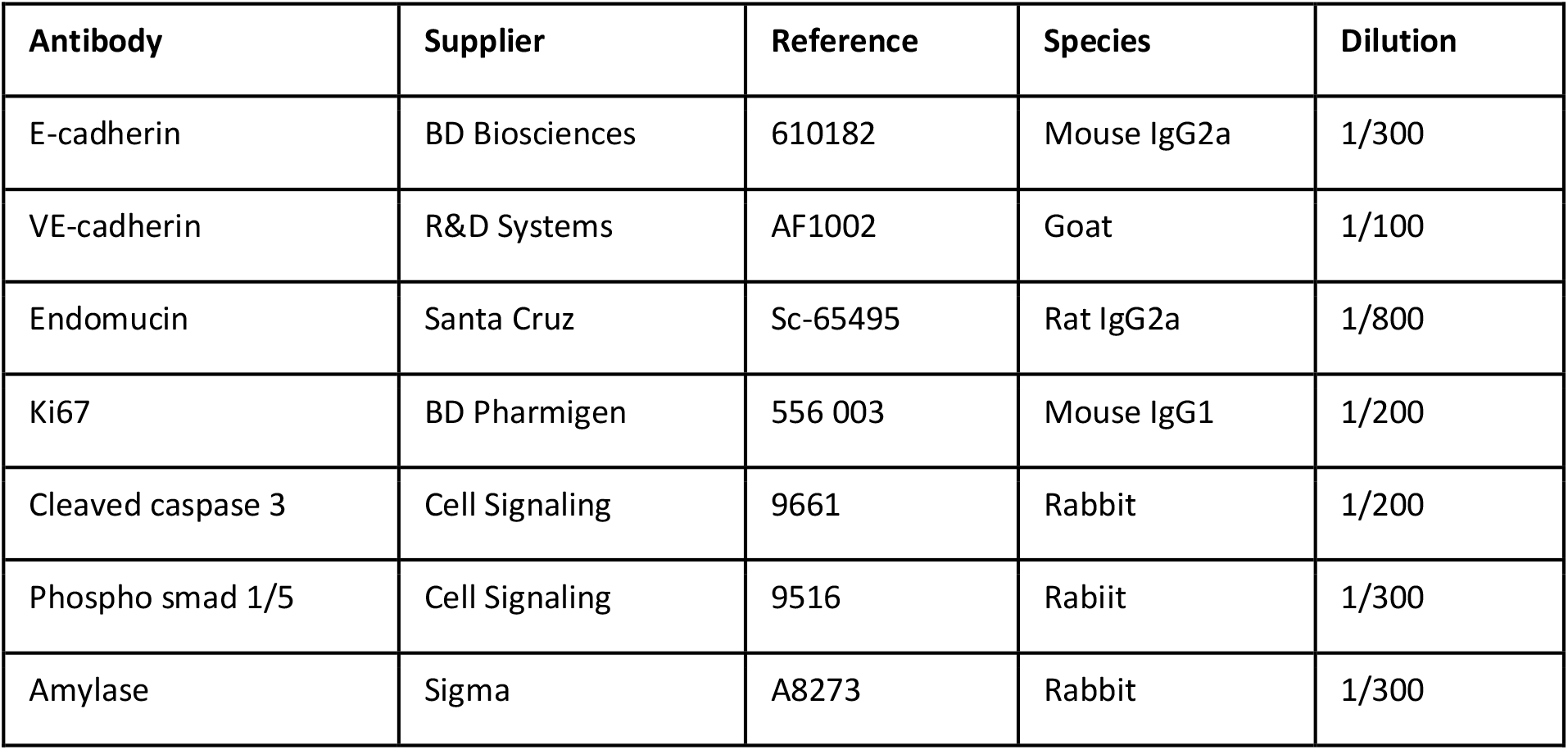
Antibodies.

| Antibody | Supplier | Reference | Species | Dilution |
| --- | --- | --- | --- | --- |
| E-cadherin | BD Biosciences | 610182 | Mouse IgG2a | 1/300 |
| VE-cadherin | R&D Systems | AF1002 | Goat | 1/100 |
| Endomucin | Santa Cruz | Sc-65495 | Rat IgG2a | 1/800 |
| Ki67 | BD Pharmigen | 556 003 | Mouse IgG1 | 1/200 |
| Cleaved caspase 3 | Cell Signaling | 9661 | Rabbit | 1/200 |
| Phospho smad 1/5 | Cell Signaling | 9516 | Rabbit | 1/300 |
| Amylase | Sigma | A8273 | Rabbit | 1/300 |

Slides were mounted and images acquired on the Cell Observer Spinning Disk Confocal Microscope (Zeiss). For quantification, positive labeled surface for Hoechst, VE-cadherin, amylase, cleaved caspase 3 or phospho-Smad 1/5 stainings, were measured with the open access image analysis Image J software, as Pearson’s correlation coefficient for colocalization between VE-cadherin and cleaved caspase 3 pixels.

### RT-qPCR

Total RNA was collected from pancreatic dorsal buds, different organs, or pancreatic explants using TRIzol reagent (Thermo Scientific) and phenol/chloroform extraction, as described[57]. Reverse transcription was performed on the total amount of extracted RNA for explants or 500 ng for non-cultured tissue samples, with random hexamer primers and the M-MLV Reverse Transcriptase (Invitrogen). Real-time quantitative PCR on cDNA was realized with the KAPA SYBR Fast qPCR kit (Sopachem) according to the manufacturer’s instructions. Primers sequences are listed in Table 3. Data were analyzed according to the Livak method (ΔΔCT) and represented as log2 fold change of the mRNA, relative to the expression of the geometric mean of the reference genes *Actb* and *Rpl27* and then compared to the control condition.

**Table 3:**
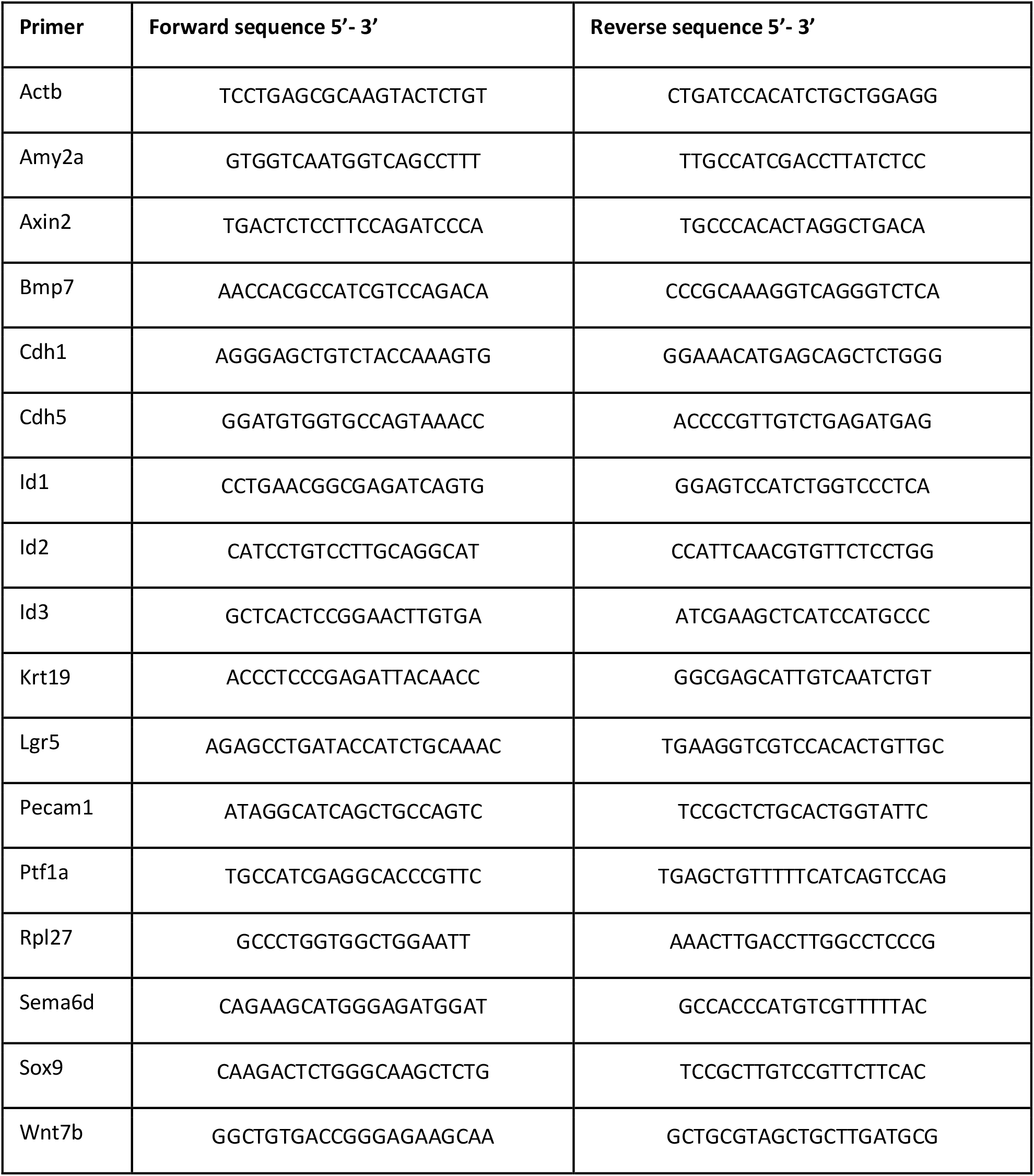
Primers.

### Statistical analysis for biological validation

For RT-qPCR results, each symbol on dot plots represents one embryo or one explant (from different litters for the same condition), and the mean +/- SEM is represented by histograms or lines (n=3 to 9). Note that for experiments on cultured explants, the untreated control condition was set to 1 (log2(1)=0) for each independent experiment. For image quantification, histograms represent the mean +/- SEM of all values across the different experiments (between 8 and 58 images representative of n=3 to 7). Parametric statistical tests were realized: paired or unpaired t-test for comparison of two conditions, and One-way ANOVA for more conditions (comparison to control condition; E11.5, pancreas or untreated). Differences were considered statistically significant when p<0.05 and illustrated; * stands for p<0.05, ** for p<0.01, *** for p<0.005 and **** for p<0.001.

## Supporting information

supplemental material-Moulis

## ACKNOWLEDGEMENTS

We (FMS, DI, CEP) acknowledge the support of the European Union’s Horizon 2020 research and innovation program Pan3DP FET Open [grant Number 800981]. MM was supported by Pan3DP, LGl holds a fellowship from the Fonds pour la formation à la Recherche dans l’Industrie et l’Agriculture (FRIA, Belgium), CV from the Fonds de la Recherche Scientifique (FRS-FNRS, Belgium) and DT is Research Associate form the Fonds de la Recherche Scientifique (FRS-FNRS, Belgium).

## AUTHOR CONTRIBUTIONS

MM performed the morphological and molecular studies, with the help of LGl. SVMR performed the bioinformatic and interactomic analyses; ND, CV, LGa, DT and PH participated in preliminary bioinformatic, cell sorting analysis or discussion. MM and CEP conceived and designed the project. FS, DI and CEP supervised the project. MM, SVMR, DI and CEP wrote the manuscript. All authors reviewed and approved the final version of the manuscript.

## DATA AVAILABILITY STATEMENT

Materials, data and associated protocols are available.

## ADDITIONAL STATEMENT

The authors declare no competing financial or non-financial interest.

