## supplemental material-Moulis for "Identification and implication of tissue-enriched ligands in epithelial-endothelial crosstalk during pancreas development"

**Table S1: List of cell population marker genes**

| Cell types | Markers | References |
|---|---|---|
| Epithelial general | Cdh1 | [1] |
| Epithelial general | Cldn6 | [2] |
| Epithelial ductal | Spp1 | [3] |
| Epithelial ductal | Sox9 | [4] |
| Epithelial acinar | Ptf1a | [5] |
| Epithelial acinar | Amy2b | [6] |
| Epithelial endocrine | Ins1 | [6] |
| Epithelial endocrine | Gcg | [6] |
| Epithelial endocrine | Pax4 | [7] |
| Epithelial endocrine | Insm1 | [8] |
| Mesenchyme general | Col1a1 | [9] |
| Mesothelial | Wt1 | [10] |
| Mesothelial | Upk3b | [11] |
| Mesenchyme | Rgs5 | [12] |
| Mesenchyme | Des | [13] |
| Neurons | Dlx2 | [14] |
| Neurons | Syt1 | [15] |
| Immune | Itgam | [16] |
| Immune | Cd53 | [17] |
| Endothelial | Cdh5 | [18] |
| Endothelial | Kdr | [19] |
| Erythrocyte | Gypa | [20] |
| Erythrocyte | Slc4a1 | [21] |
| Tip | Dll4 | [22] |
| Tip | Cd34 | [23] |
| Arterial | Sox17 | [24] |
| Arterial | Gja5 | [25] |
| Arterial | Efnb2 | [26] |
| Stalk | Vwf | [27] |
| Stalk | Cd36 | [28] |
| Venous | Nr2f2 | [29] |
| Venous | Emcn | [30] |
| Venous | Ephb4 | [31] |
| Lymphatic | Prox1 | [32] |
| Lymphatic | Flt4 | [33] |

**Table S2: Number of detected potential interactions between the sender and target populations**

| | | Target populations | | | | | |
|---|---|---|---|---|---|---|---|
| | | Epithelial | Mesenchyme | Endothelial | Immune | Neurons | Erythrocyte |
| Sender populations | Epithelial | 1582 | 1659 | 1730 | 1628 | 1567 | 1603 |
| | Mesenchyme | 2470 | 2614 | 2739 | 2585 | 2487 | 2511 |
| | Endothelial | 533 | 571 | 589 | 563 | 536 | 542 |
| | Immune | 1387 | 1440 | 1572 | 1551 | 1376 | 1413 |
| | Neurons | 547 | 568 | 594 | 555 | 542 | 553 |
| | Erythrocyte | 0 | 0 | 0 | 0 | 0 | 0 |

**Figure S1**

A

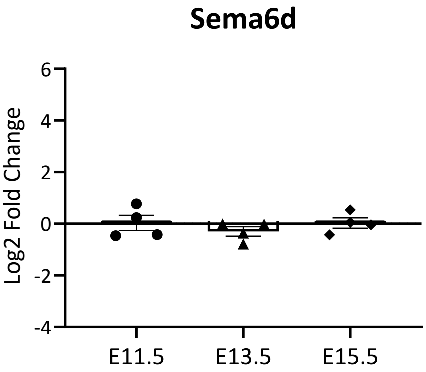


B

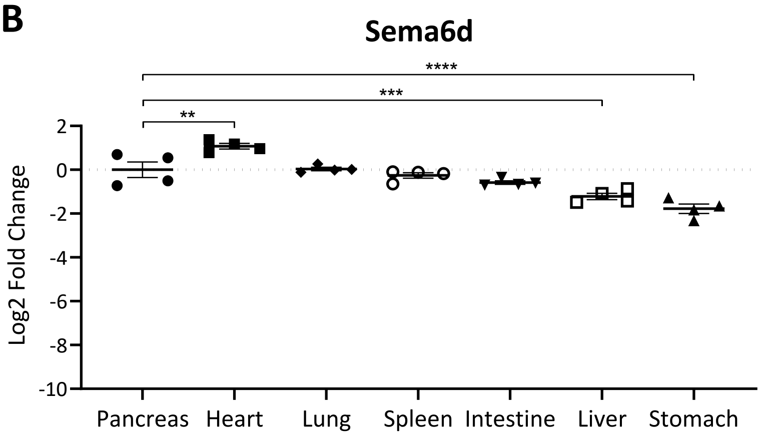


**Figure S1: Expression of Sema6d during pancreas organogenesis and in different organs. (A)** RT-qPCR analysis of Sema6d expression at three stages (E11.5 - E13.5 - E15.5) of pancreas development (n=4). Sema6d expression was stable from E11.5 to E15.5. **(B)** Comparative analysis of the expression of Sema6d by RT-qPCR in the pancreas and other organs at E15.5 (n=4), reveals similar expression levels. Actb and Rpl27 were used as housekeeping genes (ΔCt) and results are presented as log2 fold change (log2 (2-ΔΔCt)), as compared to E11.5 (A) or to pancreatic tissue (B). ** p<0.005, *** p<0.0005, **** p<0.0001.

**Figure S2**

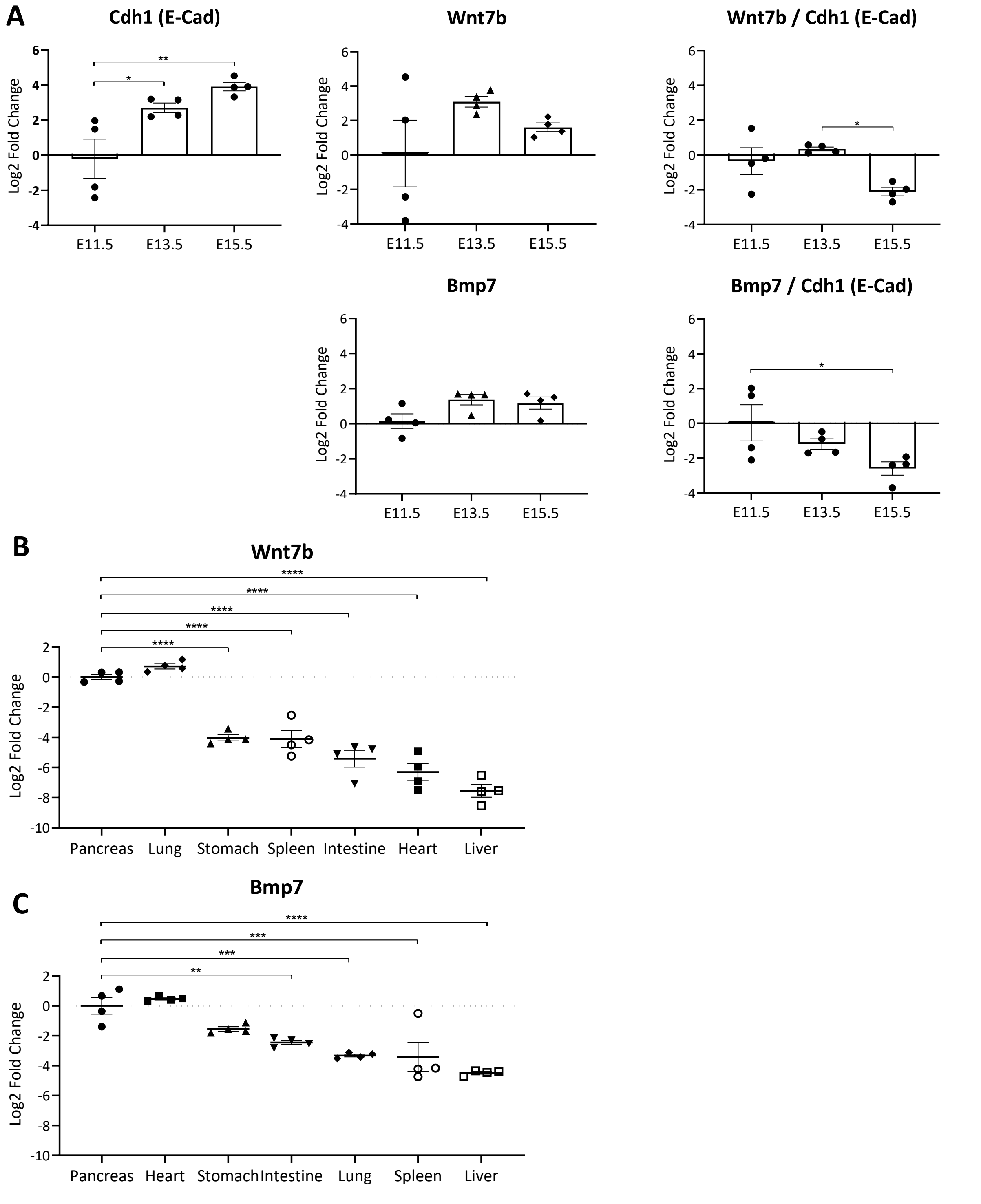


**Figure S2: Expression of Wnt7b and BMP7 during pancreas organogenesis and in different organs. (A)** RT-qPCR analysis of Cdh1, Wnt7b and Bmp7 at three stages (E11.5 - E13.5 - E15.5) of pancreas development (n=4). Increased expression of Cdh1 from E11.5 to E15.5, explains the decreased expression of Wnt7b and Bmp7 upon normalization on Cdh1 (right graphs). Comparative analysis of the expression of Wnt7b **(B)** and Bmp7 **(C)** by RT-qPCR in the pancreas and other organs at E15.5 (n=4), shows higher expression of these two transcripts in the pancreas. Actb and Rpl27 were used as housekeeping genes (ΔCt) and results are presented in log2 fold change (log2 (2-ΔΔCt)), as compared to E11.5 (A) or to pancreatic tissue (B,C). * $p<0.05$, ** $p<0.005$, *** $p<0.0005$, **** $p<0.0001$.

### Figure S3

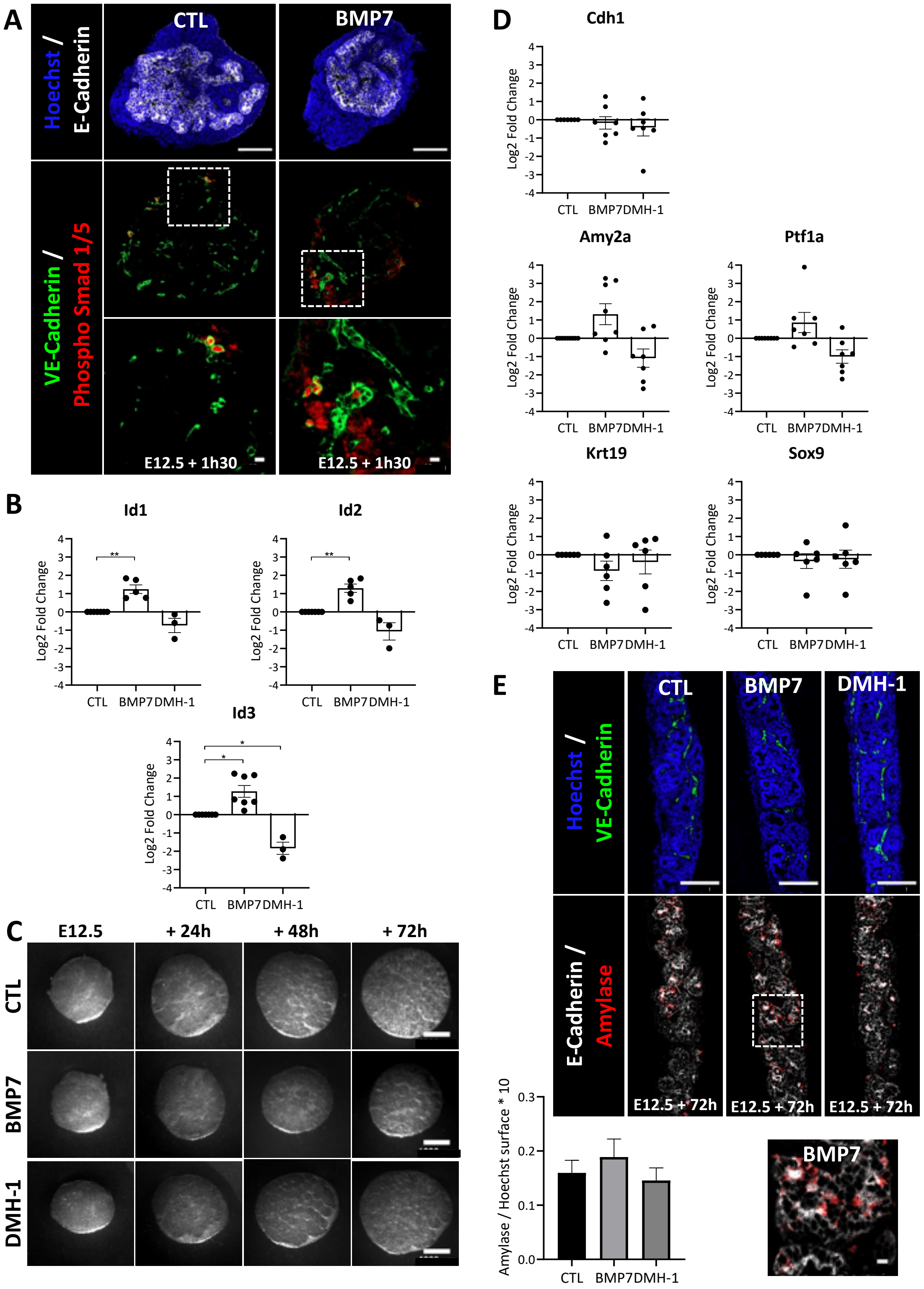


**Figure S3: Bmp7 can directly signal to endothelial cells without severely affecting the epithelial compartment. (A)** Immunolabeling of phospho Smad 1/5+ (red), epithelial E-Cad+ (white) and endothelial VE-Cad+ (green) cells, with Hoechst nuclei counterstaining (blue) on pancreatic explants treated with BMP7 for 90 minutes (BMP7, 400 ng/mL), or left untreated (CTL). Regions delineated by dashed lines are magnified below, and illustrate increased phospho Smad 1/5 positive cells (endothelial and non-endothelial) in BMP7-treated explants. Scale bar : 100 µm and 10 µm (magnification). **(B)** RT-qPCR analysis of Bmp target genes Id1, Id2 and Id3 in primary endothelial cells, normalized to Actb and Rpl27, and presented in log2 fold change, in BMP7 (400 ng/mL) or DMH-1 (3 μM) treated cells, as compared to untreated control cells cultured for 48h (n=3-7). Expression of Id genes was upregulated with BMP7 and downregulated with DMH-1. **(C)** Phase contrast images of pancreatic explants at different time points (24h - 48h - 72h) of culture without (CTL) or with BMP7 (400 ng/mL) or DMH-1 (3 μM). Besides the reduced size of BMP7 explants, treatments did not affect epithelial morphogenesis. Scale bars: 100 μm. **(D)** RT-qPCR analysis of general (Cdh1), tip (Amy2a and Ptf1a) and trunk (Krt19 and Sox9) epithelial markers normalized to Actb and Rpl27, and presented as log2 fold change, in BMP7- and DMH-1-treated explants as compared to controls cultured for 48h (n=6-9). No significant changes were measured, but a trend to an increase of tip markers was noticed. **(E)** Immunolabeling of amylase+ (red), epithelial E-Cad+ (white) and endothelial VE-Cad+ (green) cells, with Hoechst nuclei counterstaining (blue). Region delineated by dashed lines is magnified below. Quantification of the amylase+ surface reported to total Hoechst+ surface of control, BMP7- and DMH-1-treated explants cultured for 72h (n=6), revealed a slight but not significant increase of the acinar surface in BMP7 cultures. Scale bars: 100 μm and 10 μm (magnification). One-way ANOVA (comparison to CTL): * $p<0.05$, ** $p<0.005$.
